# mTOR pathway inhibition stimulates pharmacologically induced nonsense suppression

**DOI:** 10.1101/2022.08.31.506003

**Authors:** Amnon Wittenstein, Michal Caspi, Ido Rippin, Orna Elroy-Stein, Hagit Eldar-Finkelmn, Sven Thoms, Rina Rosin-Arbesfeld

## Abstract

A large number of human genetic diseases result from premature termination codons (PTCs) caused by splicing defects, insertions, deletions or point mutations also termed nonsense mutations. Nonsense mutations are the source of various genetic diseases, ranging from rare neuro-metabolic disorders to relatively common inheritable cancer syndromes and muscular dystrophies. Over the years, a wide spectrum of studies has shown that certain antibiotics and other synthetic molecules can act as nonsense mutation suppressors, by inducing readthrough of the stop-codon leading to the expression of a full-length protein. Unfortunately, most readthrough-inducing agents have limited effects and are toxic. Thus, efforts are made to improve the clinical outcome of nonsense mutation suppressors.

Here we show that the mTOR pathway is involved in antibiotic-mediated readthrough of nonsense mutations at the level of protein translation initiation. We demonstrate that inhibition of the mTOR translation-initiation-controlling eIF4E branch induces antibiotic-mediated nonsense mutation readthrough, paving the way to the development of a novel therapeutic strategy for enhancing the restoration of these disease-causing mutated transcripts.

## INTRODUCTION

Nonsense mutations are single nucleotide substitutions in the coding regions that result in premature termination codon (PTCs) leading to the production of truncated, non-functional proteins. It is well established that a wide range of human syndromes is caused by nonsense mutations in important genes [1], and a meta-analysis, based on the human gene mutation databases, showed that nonsense mutations are responsible for approximately 11% of all the gene aberrations associated with inheritable diseases [2]. Among these are disorders such as cystic fibrosis (CF), Duchenne muscular dystrophy (DMD), Usher syndrome, Rett syndrome, Ataxia-telangiectasia (A-T), hemophilia A and B (HA and HB), spinal muscular atrophy (SMA) and colorectal cancer (CRC) [3-6]. It is also known that induced production of a full-length protein, even if only to a limited extent, may be functionally and therapeutically significant. This is especially relevant for disorders where protein expression is essentially absent, such as in CF, SMA and A-T [2].

Aberrant mRNA transcripts are usually degraded by the nonsense-mediated decay (NMD) mechanism. However, some mutated mRNAs are NMD-resistant, allowing the expression of mostly non-functional, truncated protein products by causing the ribosome to release the premature peptide [7, 8]. It has been known for some years now, that certain compounds can stimulate the ribosome to misinterpret the PTC as a sense codon and thereby restore the production of the full-length protein product (reviewed in [9]). Genetic and biochemical studies have shown that nonsense mutation readthrough agents act by binding a specific site in rRNA and as a result, the ribosome introduces an amino acid instead of releasing the mRNA chain. Although little is known about the exact nature of the amino acid inserted or the precise readthrough mechanism, translation through the premature termination codon, may result in expression of a full-length protein [10]. The nucleotide context of the PTC, the position of the mutation in the mRNA and the availability of specific tRNAs and translational release factors have all been shown to play crucial roles in readthrough efficiency [11, 12]. Aminoglycoside antibiotics (AAGs) were the first drugs studied for their capacity to induce PTC-readthrough

[7]. It was shown that these compounds enable the mis-incorporation of near-cognate tRNA (nc-tRNA) at the A-site of the ribosome. Nc-tRNAs have anticodons that are complementary to only two of the three positions of a nonsense codon in mRNA [13, 14], so the insertion of an nc-tRNA overcomes the PTC and permits the translation of the full-length protein. Recent evidence indicates that AAGs function by a mechanism that competes with translation termination [15]. However, the lack of specificity, modest readthrough effects and toxicity of AAGs lead to the search for more efficient compounds [4, 16-20]. Several compounds were identified, that could increase protein production in several cell culture and animal disease models, but the readthrough levels were usually low, typically achieving no more than 5% of wild-type protein expression [21]. Importantly, in some cases, such as in lysosomal storage disease, even 1% of normal protein function may restore a near-normal or clinically less severe phenotype [22, 23], this threshold is disease and gene dependent as for cystic fibrosis, it has been shown that 10 to 35% of CFTR activity might be needed to significantly alleviate pulmonary morbidity [24] and in DMD – 1-30% of the full-length dystrophin protein is needed [9].

Following numerous reports indicating that nonsense mutation readthrough could promote only a small percentage of the full-length protein normal expression levels, Baradaran-Heravi et al. have demonstrated that readthrough activity can be chemically potentiated [21]. Other studies suggest that under certain conditions, nonsense mutation readthrough may be enhanced either by ribosomal pausing during elongation, or repression of the NMD system [25, 26].

CRC, one of the most common cancer types, can arise from a nonsense mutation in key tumor suppressor genes. CRCs progress from adenomas, which are dysplastic but non-malignant precursor lesions in the colon. Progression to carcinoma occurs through the accumulation of multiple somatic mutations, ultimately leading to malignant transformation and the formation of invasive cancer [27]. One of the most critical genes mutated in the progression of CRC is the *adenomatous polyposis coli* (APC) tumor suppressor gene [28, 29].

Since APC mutations are detected very early in the adenoma-carcinoma sequence, the APC protein is thought to act as a gatekeeper of colorectal carcinogenesis and functional loss of APC appears to be a prerequisite for malignancy development [30]. APC is a large (310 kDa) protein that has many well-characterized functional domains, including an oligomerization and an armadillo region in the N-terminus, a number of 15-and 20-amino acid β-catenin binding repeats in its central domain, and a C-terminus that contains binding sites for EB1 and the human disc large (DLG) protein. APC is a multifunctional tumor suppressor gene that is mutated in approximately 80% of both sporadic and hereditary CRC syndrome tumors [29-31]. The primary function of APC has been attributed to the negative regulation of the canonical Wnt signaling pathway through proteasomal degradation of β-catenin. In addition, APC functions beyond Wnt pathway regulation and is involved in cellular processes related to cell cycle control, migration, differentiation, and apoptosis, all of which, when dysregulated, might contribute to cancer [32-34]. Although both alleles are altered in APC-defective colorectal tumors, homozygous deletions of APC are relatively rare. Instead, more than 70% of APC mutations generate premature stop codons, resulting in stable truncated gene products, among which mutations at codons 1309 and 1450 are the most commonly found [35, 36]. Although APC deficiency that results from mutational loss of the APC C-terminal sequence is regarded as a critical event in the initiation of colon cancer, there is increasing evidence that APC truncations may exert dominant functions contributing to colon tumorigenesis. These include enhancement of cell migration, interference with spindle formation, and induction of chromosome instability [37-40]. Despite being a highly frequent mutational event in CRC, there are currently no therapeutics directly targeting APC truncations [41].

Regulating protein translation is crucial for controlling cell growth and proliferation and thus translational dysregulation may lead to aberrant growth and tumorigenicity (reviewed in [42]). Translational control is mediated by the 7-methyl-GTP cap structure present at the 5’ termini of eukaryotic mRNAs. The initiation factors 4G (eIF4G), 4E (eIF4E) and 4A (eIF4A) construct a protein complex (eIF4F), which binds to the cap structure and positions the ribosome near the 5’ terminus of the mRNA. eIF4E binding proteins (4E-BPs) inhibit protein translation by binding to eIF4E, thus preventing its association with eIF4G. Overall translation levels are therefore lowered when 4E-BP1 is active and this activity is thought to be regulated by mTOR-dependent phosphorylation [43]. mTOR (mammalian target of rapamycin) is a highly conserved serine/threonine kinase that plays a significant role in controlling cell growth and metabolism [44]. Through distinct protein complexes, it controls the levels of available cellular energy substrates and maintains the available amino acid pool by regulating protein translation. Dysregulation of the mTOR pathway leads to aberrant protein translation which manifests in various pathological conditions [45].

Here we show that inhibiting mTOR activity, specifically at the level of translation initiation, increases antibiotic-mediated nonsense suppression which opens new avenues in understanding the mechanism of ribosomal readthrough.

## MATERIALS and METHODS

### Cell culture

WT and TSC^-/-^ MEFs, APC 1450X cells [46] and human colon carcinoma cell lines (except DU4475) were cultured in Dulbecco’s modified Eagle’s medium (DMEM) supplemented with 10% fetal calf serum (FCS) and 100 U/ml penicillin-streptomycin. DU4475 were cultured in Roswell Park Memorial Institute (RPMI) 1640 Medium with 10% FCS and 100 U/ml penicillin-streptomycin. Cells were kept in a humidified 5% CO_2_ atmosphere at 37°C. All the following cell lines were from ATCC: COLO 320 - ATCC CCL-220, SW403 - ATCC CCL-230, SW620 - ATCC CCL-227, SW837 - ATCC CCL - 235, DU4475 - ATCC HTB-123, HCT116 – ATCC CCL-247 and SW1417 - ATCC CCL-238. SW48 cell line was a kind gift from Prof. Uri ben David. MEF cells deficient in TSC1/2 (MEF TSC1/2^−/−^) and matched MEF cells were generously provided by Dr. Kwiatkowski (Harvard Medical School, Boston, MA, USA) [47].

### Antibodies and reagents

The following antibodies and reagents were used: Anti-GFP (mouse monoclonal; Santa Cruz; sc-9996, 1:750), anti-APC (rabbit polyclonal; Santa Cruz; sc-7930, 1:500), anti-active β-catenin (rabbit polyclonal; Cell Signaling Technology; D2U8Y, 1:2000), anti-tubulin (mouse monoclonal; Sigma; T6199, 1:10,000), Anti-pS6K1 (rabbit polyclonal; Cell Signaling Technology; #9205, 1:1000), Anti-S6K1 (rabbit polyclonal; Cell Signaling Technology; #9202, 1:1000), Anti-4EBP-1 (rabbit polyclonal, Abcam, ab2606, 1:1000), Anti-pS6 rabbit polyclonal; Cell Signaling Technology; #2211, 1:1000), anti-mouse and anti-rabbit-HRP (Jackson Laboratories, 1:10,000), Gentamicin sulfate (Biological Industries,03-035) and G418 sulfate (Mercury-ltd, CAS 108321-42-2), Rapamycin (AdooQ BioScience, A10782) in DMSO, Torin-1 (Caymanchem, 10997) in DMSO, PF-4708671 (AdooQ BioScience, A11755) in DDW and 4EGI-1(AdooQ BioScience, A14199) in DMSO.

### Immunofluorescence

Cells were grown on 13 mm round coverslips and treated as described. The cells were fixed for 20 min in PBS containing 4% paraformaldehyde. After three washes with PBS, the fixed cells were permeabilized with 0.1% Triton X-100 for 10 min and blocked with bovine serum albumin for 1 h. Subsequently, cells were incubated at room temperature with rabbit anti-active β-catenin and anti-Rabbit for 60 and 45 min, respectively. 4’,6-Diamidino-2-phenylindole (DAPI, Sigma 10 μg/ml) was used to stain cell nuclei. An independent script was used to quantify the RGB intensity of nuclear β-catenin.

### Western blot analysis

Cells were washed with PBS and solubilized in lysis buffer (50 mM Tris pH 7.5, 100 mM NaCl, 1% Triton X-100, 2 mM EDTA) containing a protease inhibitor cocktail (Sigma). For full-length APC detection, CRC cell lines were solubilized in 6 M urea lysis buffer (50 mM Tris pH 7.5, 120 mM NaCl, 1% NP-40, 1 mM EDTA) containing protease inhibitor cocktail. Extracts were clarified by centrifugation at 12,000×g for 15 min at 4°C. Following SDS polyacrylamide gel electrophoresis (SDS-PAGE), proteins were transferred to nitrocellulose membranes and blocked with 5% low-fat milk. Membranes were incubated with specific primary antibodies, washed with PBS containing 0.001% Tween-20 (PBST), and incubated with the appropriate horseradish peroxidase-conjugated secondary antibody. After washing in PBST, membranes were subjected to enhanced chemiluminescence detection analysis. Band intensities were quantified by Fusion-Capt analysis software.

### RNA isolation and RT-qPCR analysis

Total RNA was isolated from the cultured cells using TRI reagent (Bio-lab) and an RNA extraction kit (ZYMO) according to the manufacturer’s protocol. Total RNA (1 μg) was reverse transcribed using the iScript cDNA Synthesis Kit (Bio-Rad) according to the manufacturer’s instructions. Real-time PCR was performed using the CFX Connect Real-Time PCR Detection System (Bio-Rad) using a SYBR Green Master mix (PCR Biosystems). Actin was used as a housekeeping control. All reactions were in triplicates. The primers for the amplification of the specific cDNA sequences were:

### qPCR primers

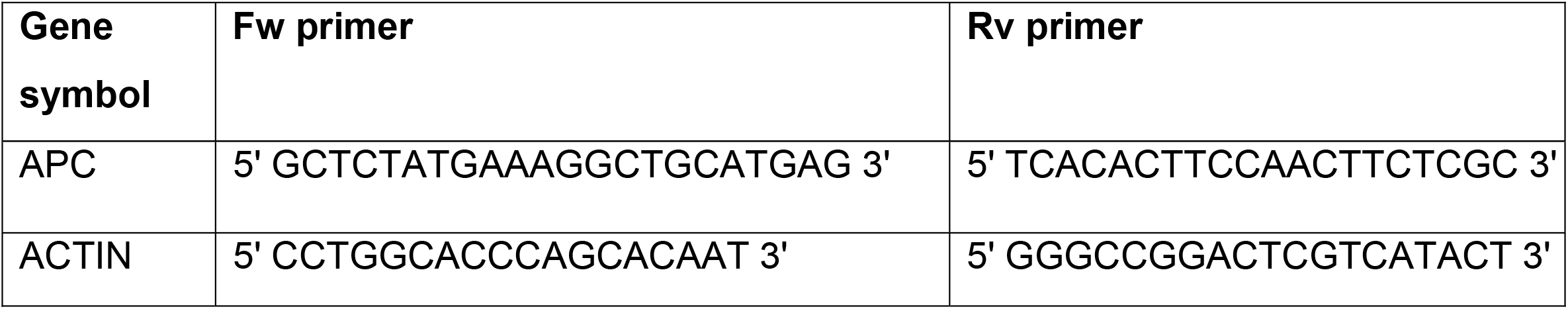

### Cell viability assay

PrestoBlue viability reagent (Thermofisher, A13261) was used according to the manufacturer’s protocol. Measurement of absorbance at 570 and 600 nm, using Epoch microplate Spectrophotometer (BioTek). All the treatments were in triplicates.

### Statistical analysis

Data were analyzed using GraphPad Prism software (version 8.0, GraphPad, La Jolla, CA) and are presented as the mean with standard deviation. Analysis of variance (ANOVA) was performed when appropriate to assess the significance of variations, using Tukey’s multiple comparisons. P values are as indicated.

## RESULTS

### Serum starvation enhances APC antibiotic-mediated nonsense-mutation readthrough in different CRC cell lines

We have previously demonstrated that stress induced by serum starvation increases antibiotic-mediated nonsense mutation readthrough in both a reporter-based cellular system and the endogenous APC gene product in the CRC cell line Colo320 [46]. To further understand the physiological mechanisms underlining this effect, we tested whether additional CRC cell lines (harboring different endogenous APC nonsense mutations) are also susceptible to serum starvation-enhanced nonsense mutation readthrough. Cells were incubated for 24h in a medium containing 10% or 1% serum supplemented with 1.5mg/ml G418. Our results show that out of the four tested cell lines – SW837, SW620, DU4475and SW1417 – only the latter was unaffected by serum starvation as reducing serum concentration to 1% did not increase antibiotic-mediated nonsense mutation readthrough activity (Fig. 1A). Similarly, other CRC cell lines with nonsense mutations in APC including SW480 and Colo320 did not respond to the treatment ([46] Fig. 1B and not shown). Neither G418 nor serum deprivation had any effect on the levels of the truncated APC (Tr-APC) form of several CRC cell lines tested (Fig.1B; Fig.S1). The specificity of antibiotic-mediated nonsense mutation readthrough was tested by comparing CRC cell lines that did not arise from an APC nonsense mutation to the Colo320 cells. Indeed, HCT116 which harbors a β-catenin deletion and SW48 which carries an APC missense mutation were not affected by G418 treatment and the levels of active β-catenin remained unchanged (Fig. 1C).

**Fig. 1.**
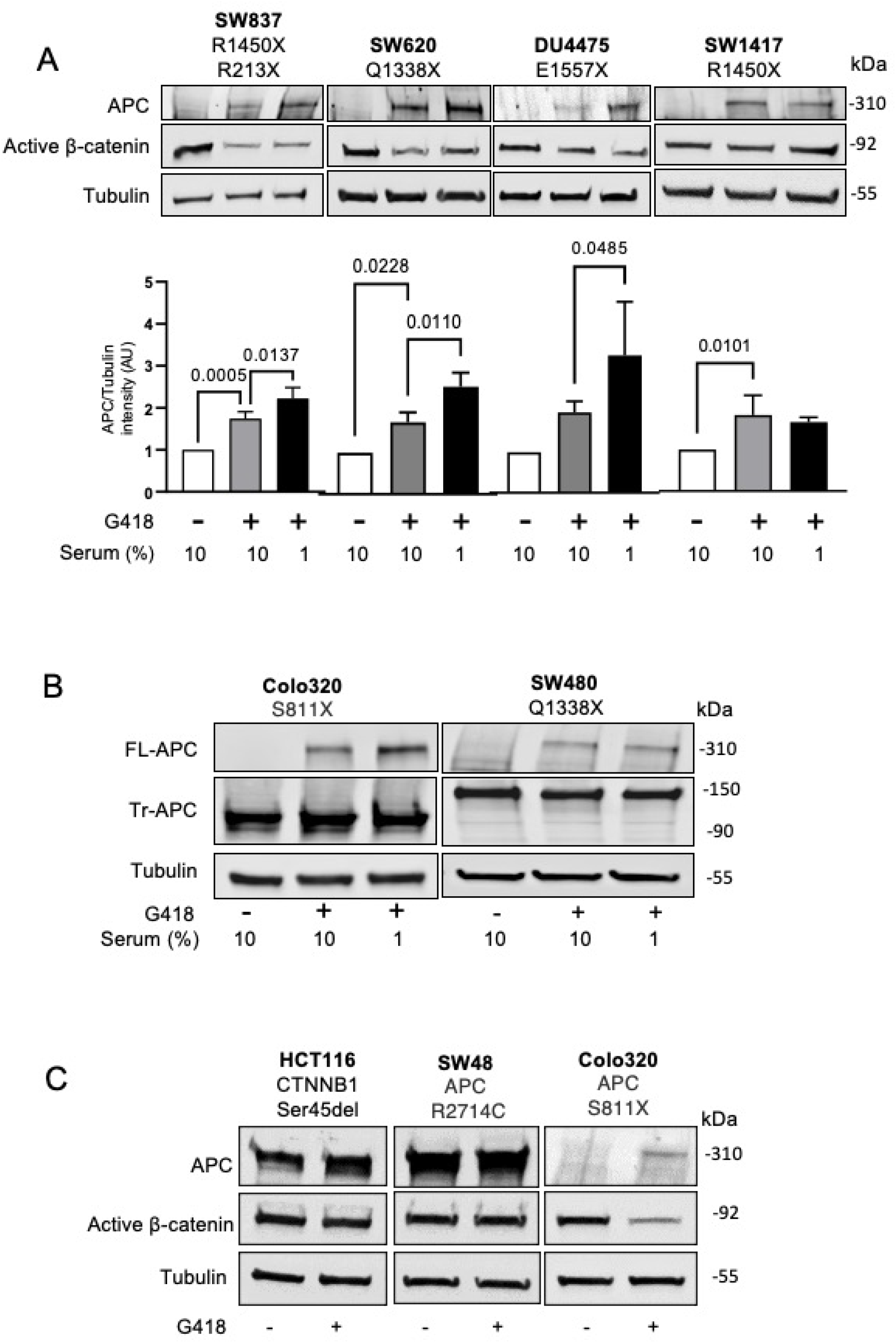
Serum starvation increased G418-mediated APC nonsense-mutation readthrough. **A**. Colon carcinoma SW837, SW620, SW1417 and breast cancer DU4475 cell lines were supplemented with 10% (NT) or 1% serum (serum starvation) as indicated and treated with 1.5 mg/ml G418 for 24h followed by western blot (WB) analysis using antibodies specific for APC, active β-catenin and tubulin. The specific mutations in each cell line are indicated. The graphs represent APC/tubulin band intensities (arbitrary units) calculated by the Fusion-Capt analysis software. Bars are mean values ± SD of 3-5 independent experiments. One-way ANOVA tests were conducted for SW837: P<0.0001, SW620: P=0.0006, SW117: P=0.0085, DU4475: P=0.0027. Tukey’s multiple comparisons scores are depicted. **B**. Colo320 and SW480 cell lines were treated as in A, followed by western blot analysis using antibodies specific for APC and tubulin. **C**. HCT116, SW48 and Colo320 cell lines were treated with 500 μg/ml G418 for 24h followed by western blot analysis using the indicated antibodies.

### The mTOR pathway may regulate antibiotic-induced nonsense mutation readthrough

To understand why some CRC cell lines (Colo320, SW837, SW620 and DU4475) respond to APC antibiotic-mediated readthrough and others (SW1417, SW403, LOVO, SW480) do not [46], we performed a two-class comparison of protein expression levels using the Broad Institute DepMap web portal (Fig. 2A). We analyzed 214 proteins that included data on all the 8 cell lines we examined, of which 8 proteins were significantly statistically different. Out of these proteins, 4EPB-1 and its three phosphorylated forms had the most statistical significance. Our analysis revealed that the eukaryotic translation initiation factor 4E (eIF4E)-binding protein 1 (4EBP-1) was highly-expressed in the cells that responded to serum starvation compared to the non-responsive cells (Fig. 2A; Fig. S2). Moreover, three additional 4EBP-1 phosphorylated isoforms also fall within the same cluster of highly-expressed proteins (Fig. 2A; Fig. S2). 4EBP-1 belongs to a family of translation repressor proteins and is a known substrate of mammalian target of the rapamycin (mTOR) signaling pathway [48]. When stimulated, mTORC1 mediates the phosphorylation of 4EBP-1 to initiate protein synthesis [49]. A schematic illustration of the main components of the mTOR signaling pathway is shown in Fig. 2B. To determine whether the mTOR cascade is involved in antibiotic-mediated nonsense mutation readthrough, we examined the effect of the tuberous sclerosis complex gene (TSC1/2) that negatively regulates mTOR signaling [50]. Immortalized TSC1/2^−/−^ mouse embryo fibroblasts (MEFs), which have a constitutively high mTOR activity [47, 50] and wild-type MEFs were transfected with the nonsense sequence readthrough-sensitive reporter plasmid GFP-BFP [24], and the effect of gentamycin (GM) on readthrough was evaluated. The GFP-BFP reporter plasmid harbors specific stop codon (and surrounding sequences) inserted between the GFP and BFP open reading frames. The upstream GFP protein serves as a control for total chimeric protein expression while the levels of GFP-BFP protein products represent the degree of stop codon readthrough activity and can be measured by Western blot analysis [24]. In the TSC1/2^−/−^ cells, no readthrough of the stop codon separating the GFP and BFP sequences was induced by the antibiotic treatment (Fig. 2C), and thus, no fusion GFP-BFP chimeric protein was detected as opposed to the wild-type cells that expressed the chimeric protein following treatment. These results suggest that the mTOR pathway may be involved in nonsense mutation antibiotic-mediated readthrough. Treating TSC1/2^−/−^ cells with rapamycin (Rap), an mTOR inhibitor, led to the increased activity of the antibiotic in inducing stop codon readthrough (Fig. 2C) further supporting this notion.

**Fig. 2.**
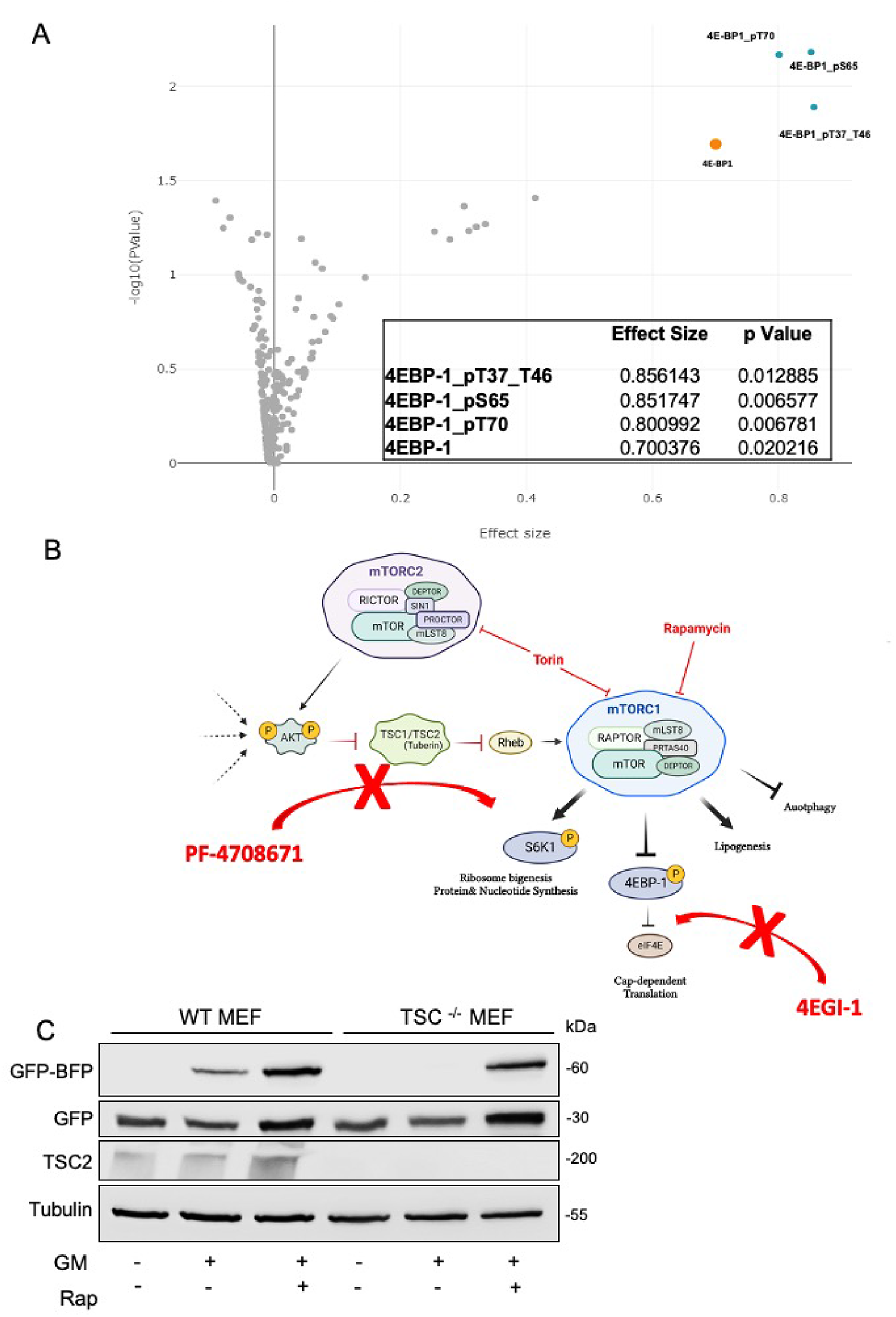
The mTOR pathway may regulate antibiotic-induced nonsense mutation readthrough. **A**. Volcano plot. Two-class comparison of protein expression levels between cell lines in which serum starvation increased APC readthrough (Colo320, SW620, SW837 and DU4475 - *in-group*) and non-responsive cell lines (SW403, SW480, SW1417 and LOVO - *out-group*) was conducted using the Broad Institute DepMap web portal (all the proteins available for these cells lines, in this database) between these two groups. **B**. A schematic illustration of the mTOR pathway and the inhibitors that were used in this study (in red). Created with BioRender.com. **C**. WT MEF and TSC ^-/-^ MEF cells were seeded in a 24-well plate before transfection. The MEFs were transiently transfected with the GFP-Stop-BFP construct (S1278X) and treated with 1.5 mg/ml gentamicin (GM) or 1 μM Rapamycin (Rap) for 24h followed by WB using the indicated antibodies.

### Torin-1 increases antibiotic-mediated nonsense codon readthrough

Torin-1 is a synthetic mTOR inhibitor that blocks ATP-binding to mTOR and thus inactivates both mTORC1 and mTORC2 [51]. Inhibition of mTOR blocks the phosphorylation of S6K and 4EBP-1 [48]. Cells stably expressing the GFP-BFP reporter plasmid (APC R1450X) were treated with Torin-1, in the presence of readthrough-inducing antibiotic gentamicin (Fig. 3A). The results show that inhibiting mTOR by Torin-1 leads to enhanced antibiotic-mediated stop codon readthrough (Fig. 3A). We then examined the effect of mTOR inhibition on antibiotic-mediated endogenous APC readthrough in the CRC cell lines Colo320, in which serum starvation increased APC readthrough (Fig. 1B) and SW403 where serum starvation did not enhance readthrough. To avoid toxicity antibiotic concentration was reduced to 500 μg/ml, (in which cell survival was 60-100%; Fig. S3). Both cell lines exhibit relatively high levels of APC restoration following AAGs treatment, however, the response to Torin-1 was similar to that of the serum starvation effect, as it increased APC readthrough in Colo320 (∼2-fold increase; Fig. 3B), but not in SW403 cells (not significant increase; Fig. 3C). Interestingly, in both cell ines, treatment esulted in a comparable decreased expression of β-catenin. Additional cell lines that responded to serum starvation also responded to the Torin-1 treatment (Fig. S4).

**Fig. 3.**
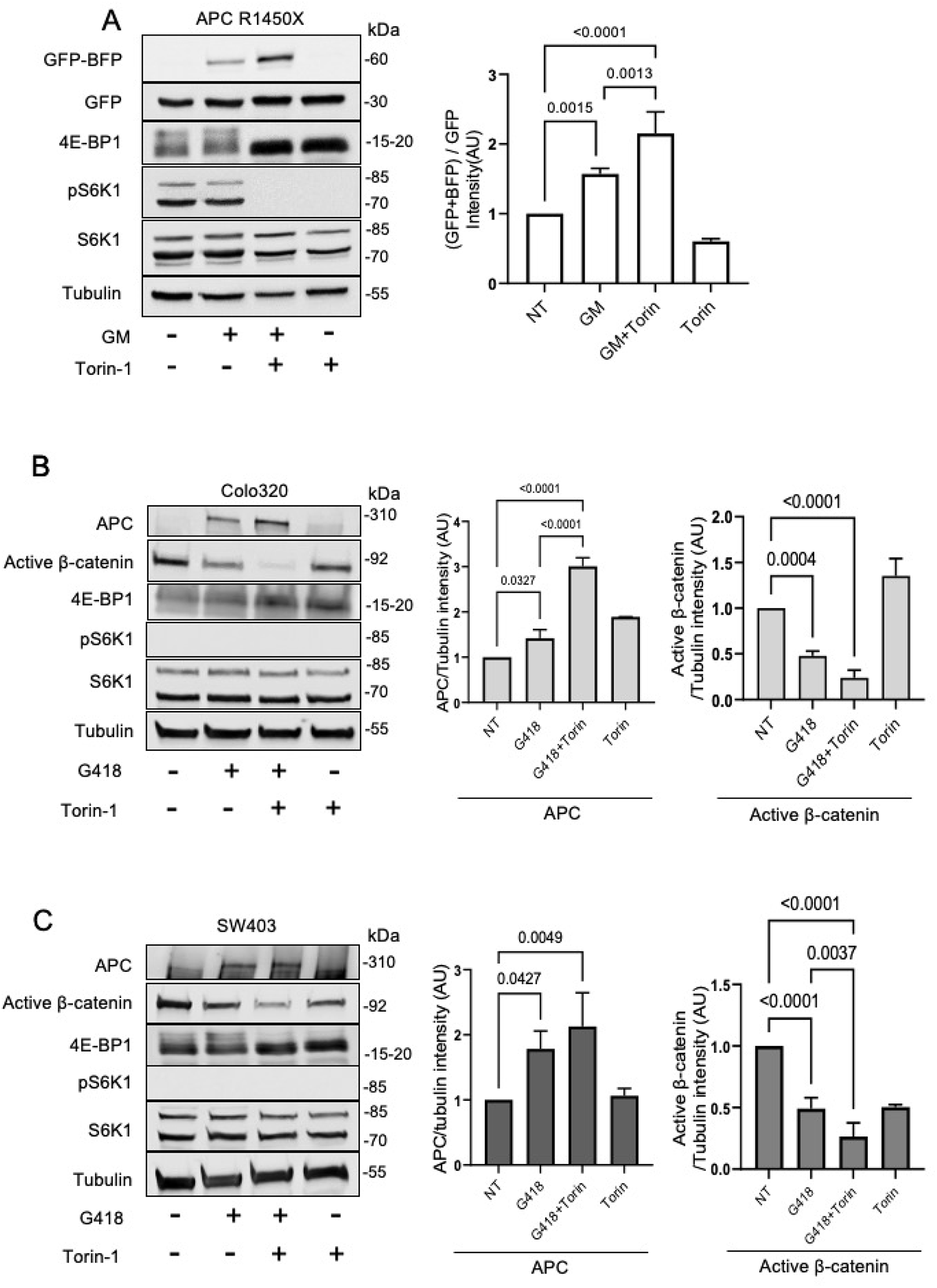
Torin-1 increases antibiotic-mediated nonsense codon readthrough. **A**. The APC R1450X reporter cell line was treated for 24h with 500 μg/ml GM and/or 500 nM Torin-1 followed by WB. The graphs show the relative GFP -BFP band intensity (normalized to GFP band intensity). Bars represent the mean values ± SD from 5 independent experiments. P<0.0001. **B-C**. Colo320 (B) and SW403 (C) were treated for 24h with 500 μg/ml G418 and/or 500 nM Torin-1 followed by WB analysis for the indicated proteins. Graphs representation of the APC/tubulin or Active β-catenin/tubulin band intensities (arbitrary units), calculated by the Fusion-Capt analysis software. The bars represent the mean values ± SD from 4 independent experiments. Colo320: P<0.0001, SW403: APC P=0.0023, Active β-catenin P<0.0001. Tukey’s multiple comparisons scores are depicted

As Torin-1 inhibits both mTORC1 and mTORC2 (Fig. 2B), we speculated that it may have additional effects on stop-codon mediated readthrough activity. We therefore tested the effects of the mTORC1 specific inhibitor Rapamycin.

### Rapamycin increases antibiotic-mediated nonsense codon readthrough

Rapamycin is a naturally occurring macrolide known to inhibit cap-dependent mRNA translation [52]. Similar to the results obtained with Torin-1, Rapamycin significantly increased antibiotic-mediated readthrough of the GFP-BFP fusion protein in APC R1450X stable cell line (Fig. 4A). In the CRC cell lines tested, the effect of rapamycin was, again, similar to the serum starvation effect, as it increased endogenous APC nonsense mutation readthrough (induced by G418) in Colo320, but not in SW403 cells (Fig. 4B-C). As expected, a decrease in active β-catenin levels was only observed in the Colo320 cell line (Fig. 4B). Interestingly, although mutated APC transcripts are relatively stable, as they often escape NMD, some increase in mRNA levels was observed in treated Colo320 cells as opposed to SW403 where mRNA transcripts were unaffected by readthrough or readthrough enhancement (Fig. S5). Next, the levels of nuclear active β-catenin following treatment were examined. In accordance with the previous results, rapamycin treatment led to reduced translocation of β-catenin into the nucleus only in Colo320 cells (Fig. 4D).

**Fig. 4.**
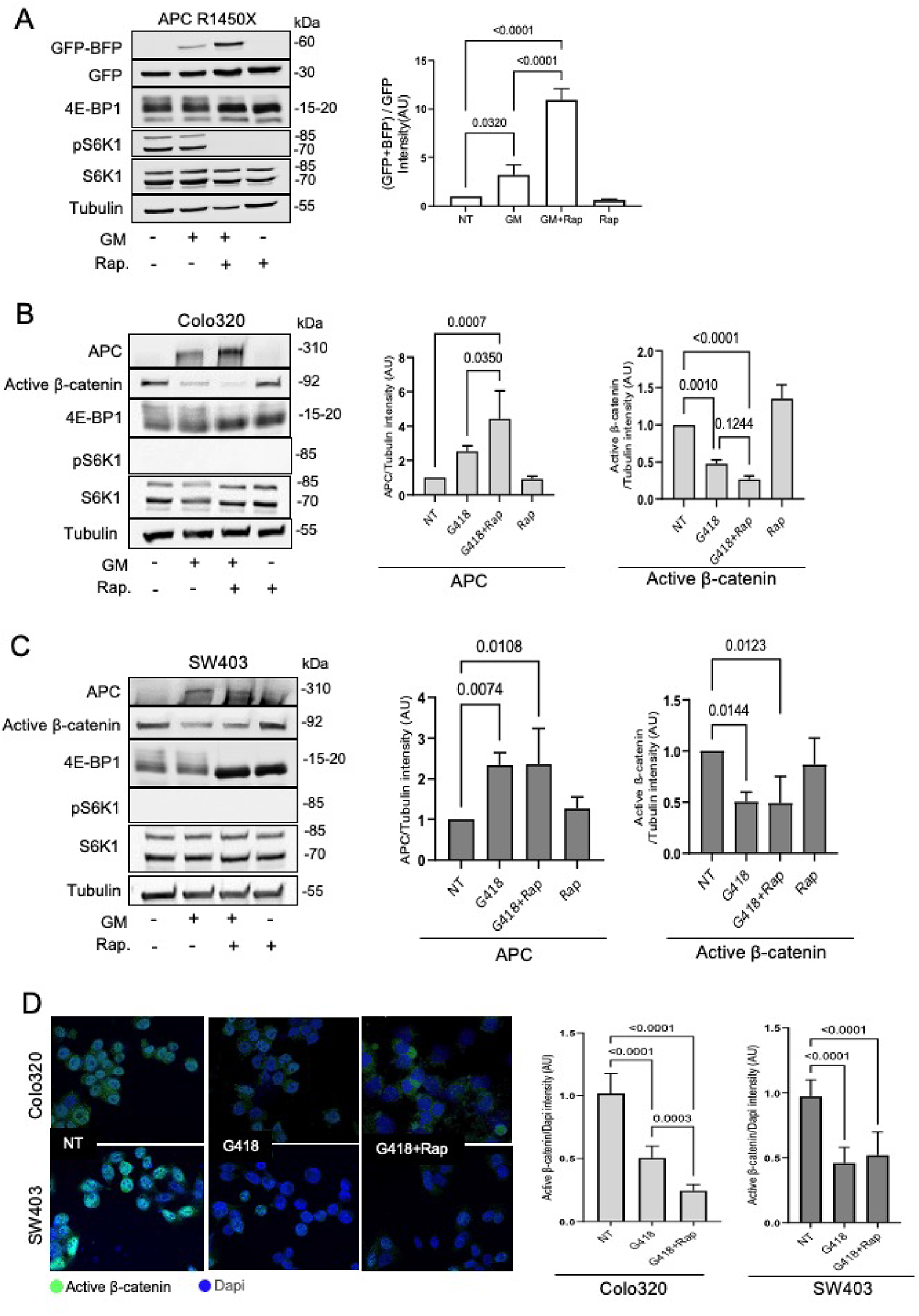
Rapamycin increases antibiotic-mediated nonsense codon readthrough. **A**. The APC-1450X reporter cell line was treated for 24h with 500 μg/ml GM or 1 μM Rap followed by WB. The graphs show the relative GFP-BFP band intensity (normalized to GFP band intensity). Bars represent the mean values ± SD from 5 independent experiments. P<0.0001. **B-C**. Colo320 (B) and SW403 (C) were treated for 24h with 500 μg/ml G418 or 1μM Rap followed by WB. Graphs represent the APC/tubulin or Active β-catenin/ tubulin band intensities (in arbitrary units), calculated by the Fusion-Capt analysis software. The bars represent the mean values ± SD from 4-6 independent experiments. Colo320: APC P= 0.0005, Active β-catenin P<0.0001, SW403: APC P= 0.0031, Active β-catenin P= 0.0048. **D**. Colo320 (one experiment) and SW403 (two independent experiments) were treated with 1.5 mg/ml G418 and 25μM Rap for 24h. The cells were then fixed and visualized by confocal microscopy. The graph represents the Active β-catenin intensity (green) normalized to DAPI (blue) in antibiotic-treated cells. An independent script was used to quantify the RGB intensity of nuclear β-catenin on ten independent fields of each sample. P<0.0001. Tukey’s multiple comparisons scores are depicted.

### Inhibition of eIF4E increases antibiotic-mediated nonsense codon readthrough

Activated mTORC1 signaling leads to the phosphorylation of several downstream components among them are S6K1 and 4EBP-1 (Fig. 2B). S6K1 phosphorylation is a known stimulus of protein synthesis [53], however, our results indicate that inhibiting S6K1 (using the S6K1 inhibitor (PF-4708671)) did not affect antibiotic-mediated nonsense suppression (Fig. 5A). Moreover, as Colo320 and SW403 do not express p-S6K1 (Fig. 3B-C, Fig. 4B-C) we concluded that S6K1 is not involved in the effects of mTOR inhibition on antibiotic mediates nonsense mutation readthrough.

We then examined the 4EBP-1 phosphorylation branch, that came up in our analysis (Fig. 2) and allows cap-dependent translation. The small-molecule inhibitor, 4EGI-1 stabilizes the eIF4E/4EBP-1 interaction and inhibits the elF4E-elF4G interaction thus inhibiting translation initiation [54]. Our data demonstrate that 4EGI-1 enhances antibiotic-induced nonsense mutation readthrough of both the GFP-BFP chimeric protein in APC R1450X stable cell line (Fig. 5B) and endogenous APC in Colo320 CRC cell lines (Fig. 5C), further supporting our hypothesis that inhibition of protein translation initiation enhances antibiotic-mediated nonsense mutation readthrough. Interestingly, similar results were observed in the SW403 CRC cell line (Fig. 5D), although this cell line falls into a group in which serum starvation did not affect readthrough levels (Fig. 2). Our analysis (Fig. 2A) indicates that in these cells the levels of phosphorylated 4E-BP1 are already low and thus cannot be further reduced by mTOR inhibition. However, it is possible that there are sufficient amounts of eIF4E and eIF4G in these cells and that their activity is affected by the 4EGI-1 inhibitor.

**Fig. 5.**
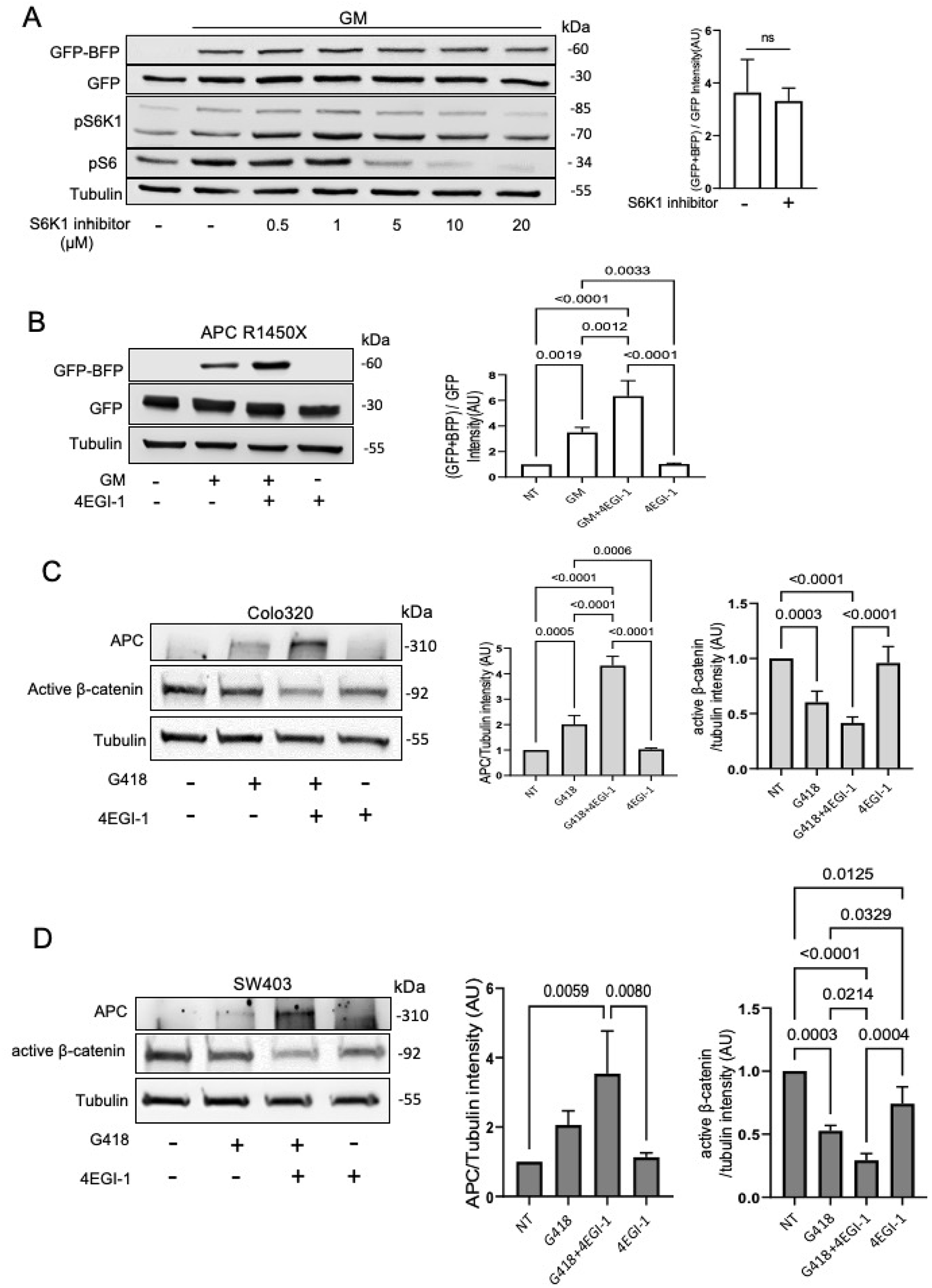
Inhibition of eIF4E increases antibiotic-mediated nonsense codon readthrough. **A**. The APC-1450X reporter cell line was treated for 24h with 500 μg/ml GM and 0.5, 1, 5, 10, or 20 µM PF-4708671 (S6K1 inhibitor) followed by WB analysis using the indicated antibodies. The graphs show the relative GFP-BFP band intensity (normalized to GFP band intensity; with or without 20 μM PF-4708671), calculated by the Fusion-Capt analysis software. The bars represent the mean values ± SD from 4 independent experiments. **B**. The APC R1450X reporter cell line was treated for 24h with 500 μg/ml Gentamicin (GM) and/or 50 μM 4EGI-1 (eIF4E/eIF4G Interaction Inhibitor). The graphs show the relative GFP-BFP band intensity (normalized to GFP band intensity) Bars represent the mean values ± SD from 5 independent experiments. P= 0.0004. **C**. Colo320 cell line was treated for 24h with 500 μg/ml G418 or 50 μM 4EGI-1. The bars represent the mean values ± SD from 4-6 independent experiments. P< 0.0001. **D**. SW403 cell line was treated for 24h with 500 μg/ml G418 or 50 μM 4EGI-1. The bars represent the mean values ± SD from 3 independent experiments. P= 0.0047 (left), P<0.0001 (right). Tukey’s multiple comparisons scores are depicted.

Taken together our results suggest that the mTOR pathway is involved in nonsense suppression control and inhibition of this pathway may have potential therapeutic roles.

## DISCUSSION

Single nucleotide substitutions in gene coding regions can change a sense codon into a nonsense or premature termination codon (PTC). The PTC-containing mRNA may be degraded by the nonsense-mediated mRNA decay (NMD) cellular surveillance pathways or translated into a truncated mostly nonfunctional protein. In both scenarios, PTCs can result in a wide range of human genetic disorders including colorectal cancer (CRC). CRC is a frequent and lethal cancer type in which nonsense mutations in at least one of the APC gene alleles account for approximately 20-30% of all cases [55-57]. Mutated APC transcripts are often NMD-resistant as most of the nonsense mutations occur in a hotspot within the last APC exon and therefore are not recognized by the exon junction complex that induces NMD [57]. As a result, many cancer cells express a truncated APC protein that is proposed to have both gain-of-function and loss-of-function activities [31]. Thus, restoring APC’s full-length function may have enormous therapeutic value [58]. The expression levels of full-length APC vary in different CRC cells harboring different mutations (such as HCT116 that expresses a mutated β-catenin but a wild-type APC form or SW48, which contains an APC missense mutation; Fig. 1C). As expected, in these cases, APC was not affected by readthrough induction. In general, suppression of nonsense mutations is a viable therapeutic strategy that is extensively studied. There are several different approaches to overcome the disease-causing phenotype of these mutations including antisense oligonucleotides, suppressor tRNAs that can read PTCs, RNA editing, or CRISPR technology [10]. However, the most commonly studied method for inducing PTC readthrough is by using AAGs [15, 59-61], macrolides [24], synthetic AAGs such as ELX-02 (NB124) [62, 63], and oxadiazole derivates such as Ataluren (PTC124) [64, 65]. Unfortunately, these compounds have limitations that include high toxicity and adverse side effects [10]. It was also shown that AAGs such as G418, cause mis-incorporation of sense-amino acids that ultimately interferes with protein synthesis [66], and thus methods that would reduce dosage or decrease treatment frequency are greatly needed.

Here we explored the possibility that PTC readthrough could be increased by modulating endogenous signaling pathways. As serum starvation induces antibiotic-mediated nonsense mutation readthrough [46] and was shown to inhibit the mTOR pathway [67], we examined the effects of mTOR inhibitors like Torin-1 and rapamycin on antibiotic-mediated readthrough. Our results show that inhibiting mTOR1 using different methods increases the PTC readthrough of both a reporter fusion protein and the endogenous APC protein. To identify the mechanisms that underlay this effect, we examined different branches of the mTOR pathway. We found that the small molecule 4EGI-1, which inhibits mTORC1 signaling and cap-dependent translation initiation (by binding to eIF4E and disrupting the eIF4E/eIF4G association) [54, 68], induces antibiotic-mediated readthrough. As CRC cells that responded to mTOR inhibition by increased PTC readthrough show high levels of 4EPB-1 (Fig. 3-4 and data not shown), we conclude that inhibiting cap-dependent protein translation initiation enhances antibiotic-mediated PTC readthrough. 4EGI-1 is of particular interest since it has anti-tumorigenic activity and reduces the growth of human cancer xenografts *in vivo* [69]. A recent study demonstrated that low concentrations of rapamycin (0.3-10nM) do not affect AAG-induced readthrough in HDQ-P1 cancer cells even though phosphorylation levels of S6K1 were decreased [66]. It was shown previously that in several cancer cell lines low concentrations of rapamycin decrease only S6K1 phosphorylation but not 4EBP-1 [70]. These observations are in line with our hypothesis that 4E-BP1 inhibition is the mediating factor in increasing AAG-induced readthrough activity.

Combining efficient nonsense suppression agents that target distinct components of the translation machinery may be a promising treatment strategy for PTC-derived diseases as it was recently shown that depletion of eRF1 induces translational pausing at PTCs and increases readthrough activity of AAGs [62, 71, 72]. Our results indicate that 4EG1-1, which affects translation initiation, enhances PTC readthrough in the presence of AAGs. AAGs exert their PTC readthrough activity by binding at the decoding center of the eukaryotic ribosome and reducing the ability of translation termination factors to accurately recognize the PTC [73, 74]. Other compounds such as the small molecules CDX5-1 [21] or the drug mefloquine [75], that do not induce readthrough when used as single agents, enhance AAG-mediated readthrough activity, possibly by targeting the translation machinery in a still unclear mechanism. It is also possible that the ability of 4EGI-1 to potentiate PTC readthrough stems from its inhibitory effect on mTORC1 signaling, which may be independent of its role in cap-dependent translation initiation [68]. As AAGs induce readthrough by reducing ribosomal proofreading during translation, a combination with agents that affect translation may lead to the development of improved readthrough-inducing compounds.

In conclusion, our findings indicate that the effects of AAGs on PTC readthrough can be increased by inhibiting the mTOR1 pathway and specifically cap-dependent translation initiation. Following up this line of research may improve our abilities to stimulate PTC readthrough for the treatment of genetic diseases caused by nonsense mutations.

## Figure Legends

**Fig. S1. Truncated APC is not affected by antibiotic-mediated nonsense mutation readthrough.** W403 and SW480 cell lines were treated with 1.5 mg/ml G418 for 24h followed by WB analysis using antibodies specific for APC and tubulin. FS= Frameshift.

**Fig. S2. Expression of 4EBP-1 in different CRC cell lines.** Two-class comparison of protein expression of 4EBP-1 and its phosphorylation derivatives between cell lines in which serum starvation increased APC readthrough (Colo320, SW620, SW837 and DU4475 – in-group) and non-responsive cells (SW403, SW480, SW1417 and LOVO – out-group) was conducted using the Broad Institute DepMap web portal (https://depmap.org/portal/download/).

**Fig. S3. Effects of treatment on cell survival.** The APC-1450X reporter cell line, Colo320 and SW403 cell lines were treated for 24h with 500 μg/ml GM (reporter cell line) or G418 (Colo320 and SW403), 500 nM Torin-1 or 1 μM Rap. PrestoBlue reagent was added to the wells and absorbance was measured after 3h incubation at 570 and 600 nm. The curves represent the mean values ± SD from 3 independent experiments.

**Fig. S4. APC restoration in different CRC cell lines.**SW620 and SW837 were treated for 24h (SW620) or 48h (SW837) with 500 μg/ml G418 and/or 500 nM Torin-1 followed by WB analysis for the indicated proteins.

**Fig. S5. APC mRNA levels in the different treatments.**Colo320 and SW403 cell lines were treated for 24h with 500 μg/ml G418 and/or 1 μM Rap. Total RNA was extracted from the treated samples, converted to cDNA, and subjected to RT-qPCR analysis. APC transcript levels were analyzed. The bars represent the mean values ± SD from 3 independent experiments. Two-way ANOVA (P=0.0012) with Tukey’s multiple comparisons test was applied - significant scores were depicted.

## Acknowledgments

The work was supported by the German-Israel Foundation (GIF) grant I-1459-412.13/2018 to RRA and ST. The authors thank Omri Kazelnik for technical advice and Julia Hofhuis for help with the figures.

